# Reannotation of the ribonucleotide reductase in a cyanophage reveals life history strategies within the virioplankton

**DOI:** 10.1101/467415

**Authors:** Amelia O. Harrison, Ryan M. Moore, Shawn W. Polson, K. Eric Wommack

## Abstract

Ribonucleotide reductases (RNRs) are ancient enzymes that catalyze the reduction of ribonucleotides to deoxyribonucleotides. They are required for virtually all cellular life and are prominent within viral genomes. RNRs share a common ancestor and must generate a protein radical for direct ribonucleotide reduction. The mechanisms by which RNRs produce radicals are diverse and divide RNRs into three major classes and several subclasses. The diversity of radical generation methods means that cellular organisms and viruses typically contain the RNR best-suited to the environmental conditions surrounding DNA replication. However, such diversity has also fostered high rates of RNR misannotation within subject sequence databases. These misannotations have resulted in incorrect translative presumptions of RNR biochemistry and have diminished the utility of this marker gene for ecological studies of viruses. We discovered a misannotation of the RNR gene within the *Prochlorococcus* phage P-SSP7 genome, which caused a chain of misannotations within commonly observed RNR genes from marine virioplankton communities. These RNRs are found in marine cyanopodo- and cyanosiphoviruses and are currently misannotated as Class II RNRs, which are O_2_-independent and require cofactor B_12_. In fact, these cyanoviral RNRs are Class I enzymes that are O_2_-dependent and may require a di-metal cofactor made of Fe, Mn, or a combination of the two metals. The discovery of an overlooked Class I β subunit in the P-SSP7 genome, together with phylogenetic analysis of the α and β subunits confirms that the RNR from P-SSP7 is a Class I RNR. Phylogenetic and conserved residue analyses also suggest that the P-SSP7 RNR may constitute a novel Class I subclass. The reannotation of the RNR clade represented by P-SSP7 means that most lytic cyanophage contain Class I RNRs, while their hosts, B_12_-producing *Synechococcus* and *Prochlorococcus*, contain Class II RNRs. By using a Class I RNR, cyanophage avoid a dependence on host-produced B_12_, a more effective strategy for a lytic virus. The discovery of a novel RNR β subunit within cyanopodoviruses also implies that some unknown viral genes may be familiar cellular genes that are too divergent for homology-based annotation methods to identify.

## 1 Introduction

Viruses are the most abundant biological entities on the planet, with an estimated 10^31^ viral particles globally (Suttle, 2005). While viruses are known to infect cellular life from all three domains, viruses largely influence ecosystems through the infection of microbial hosts. In the oceans, 10^23^ viral infections are estimated to take place every second, resulting in the mortality of approximately 20% of marine microbial biomass each day (Suttle, 2007). Cell lysis resulting from viral infection influences ocean biogeochemical cycling by returning particulate and dissolved organic matter to the water column (Jover et al., 2014; Suttle, 2005), where it may be taken up by microbial populations to fuel new growth, or exported to the deep ocean (Laber et al., 2018; Suttle, 2007). Viral predation can also influence biogeochemical cycles through the restructuring of microbial populations (Rastelli et al., 2017), metabolic reprogramming of host cells (Lindell et al., 2005; Puxty et al., 2016), and horizontal gene transfer (Lindell et al., 2004).

While the importance of viruses within marine microbial communities is now commonly accepted, the biological and ecological details of viral-host interactions that influence the transformations of nutrient elements in ecosystems are largely unknown. Attempting to reveal these details, researchers have turned to metagenomics and metatranscriptomics for assessing the genetic repertoire and biological potential of unknown microbial and viral populations (Brum et al., 2015; Coutinho et al., 2017; Moniruzzaman et al., 2017; Roux et al., 2016). Bridging the gap between genetic observations and ecosystem-level effects requires an understanding of the connections between genes and phenotypes. Among viruses infecting marine microbes, genes involved in nucleotide metabolism and viral replication are highly predictive of viral phenotype and evolutionary history (Dolja and Koonin, 2018; Iranzo et al., 2016; Kazlauskas et al., 2016).

For example, a point mutation in motif B of the family A DNA polymerase gene (*polA*) is indicative of viral life style (Chopyk et al., 2018; Schmidt et al., 2014). Another useful viral marker gene is ribonucleotide reductase (RNR). RNRs catalyze the rate-limiting step of DNA synthesis (ribonucleotide reduction) (Ahmad et al., 2012; Kolberg et al., 2004), and are therefore prominent in the genomes of lytic dsDNA phage (Dwivedi et al., 2013; Iranzo et al., 2016; Sakowski et al., 2014). They are ancient enzymes thought to have been essential in the transition from an RNA world to a DNA world (Lundin et al., 2015; Wächtershäuser, 2006) and have evolved into several classes and subclasses with diverse biochemical mechanisms and nutrient requirements (Nordlund and Reichard, 2006). Thus, the biochemical class of RNR used by a cell or virus can reflect the environmental conditions surrounding DNA replication (Cotruvo et al., 2011; Reichard, 1993; Sakowski et al., 2014).

All RNRs share a common catalytic mechanism in which a thiyl radical in the active site removes a hydrogen atom from the 3’ hydroxyl group of the ribose sugar, thereby activating the substrate (Licht et al., 1996; Logan et al., 1999; Lundin et al., 2015). The mechanism by which the thiyl radical is generated varies greatly among RNRs and provides the biochemical basis dividing the three major RNR classes (Lundin et al., 2015). Extant RNRs are also commonly divided by their reactivity with O_2_ (Reichard, 1993): Class I RNRs are O_2_-dependent; Class II RNRs are O_2_-independent; and Class III RNRs are O_2_-sensitive (Fig. 1a).

**Figure 1.**
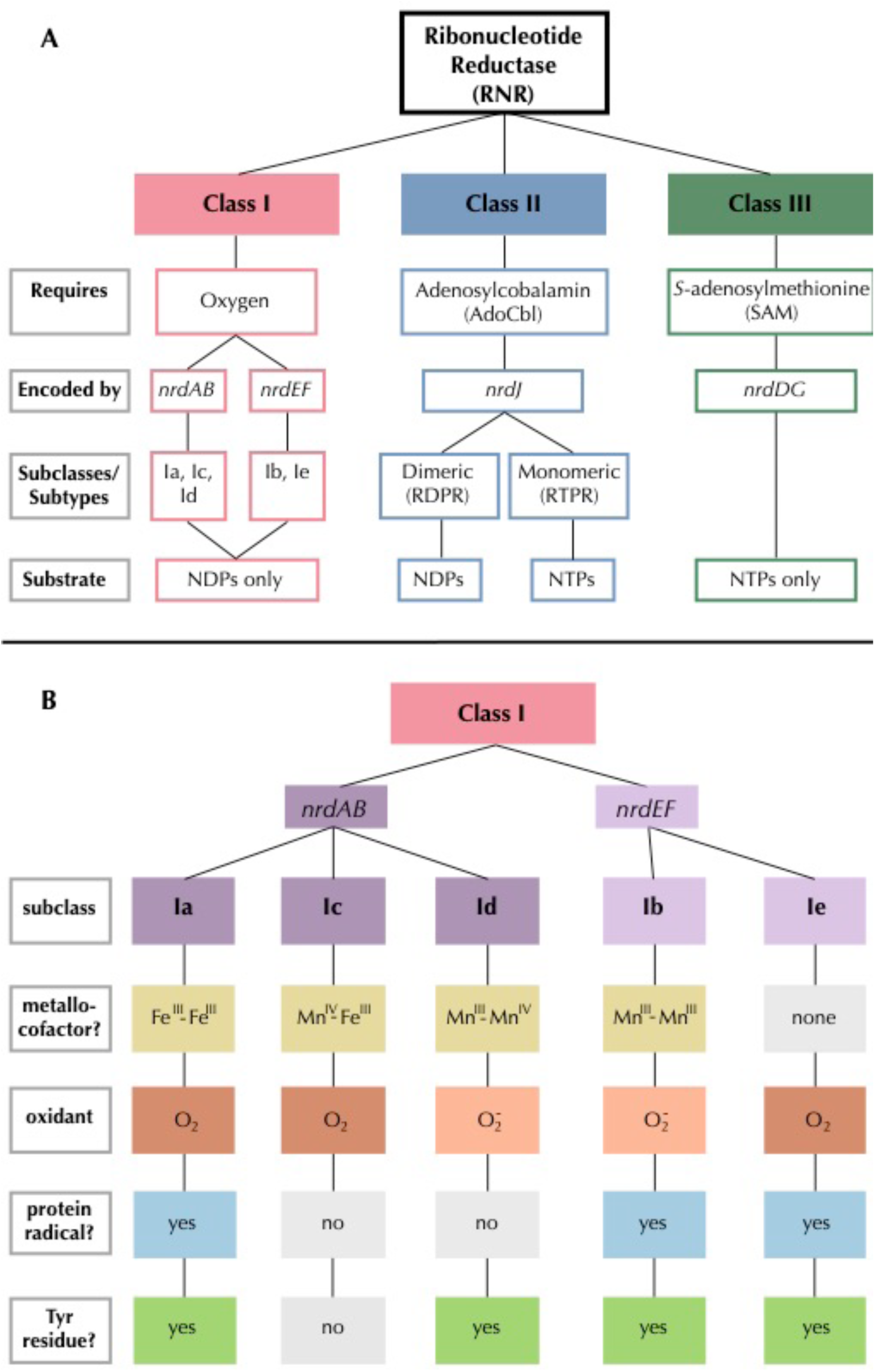
Summary of **A)** RNR class and **B)** Class I subclass divisions. Gray outlined boxes to the left indicate categories. In B, like colors indicate common traits and light gray filled boxes indicate missing traits.

Class III RNRs are the most ancient form of the enzyme and the most dissimilar of the extant types (Aravind et al., 2000; Lundin et al., 2015). They produce a radical on a small activase subunit, NrdG, before passing it to a larger catalytic subunit, NrdD (Nordlund and Reichard, 2006). The activase is a radical SAM protein, which creates a radical by cleaving *S*-adenosylmethionine using an iron-sulfur cluster (Mulliez et al., 1993). Like other glycyl radical enzymes, Class III RNRs temporarily store this radical on a glycine residue in the C-terminus of the catalytic subunit. In the presence of oxygen, the glycyl radical reacts immediately with O_2_, resulting in fragmentation and inactivation of NrdD (Eliasson et al., 1992; King and Reichard, 1995). Therefore, Class III RNRs are found only in strict or facultative anaerobes and their viruses (Fontecave et al., 2002).

Class II RNRs emerged after Class III (Lundin et al., 2015) and are the only RNRs that do not require separate subunits for radical generation and catalysis (Nordlund and Reichard, 2006). Instead, Class II RNRs are encoded by a single gene, *nrdJ*. Class II RNRs require adenosylcobalamin (AdoCbl), a form of B_12_, to produce a radical, which is then shuttled along the enzyme to the active site (Blakley and Barker, 1964; Lundin et al., 2010). There are two types of Class II RNR: monomeric and dimeric (Nordlund and Reichard, 2006). The monomeric form is commonly referred to as ribonucleotide triphosphate reductase (RTPR). Monomeric and dimeric Class II RNRs are phylogenetically distinct (Lundin et al., 2010), and it is unclear which evolved first (Lundin et al., 2015), although there is some speculation that the monomeric form arose from the duplication of a portion of the gene encoding the dimeric form (Sintchak et al., 2002).

Class I RNRs are the most recent (Lundin et al., 2015) and the most complex of the extant RNRs (Fig. 1b). Like the Class III RNR, radical generation takes place on a smaller subunit (β or R2) and is transferred to a larger catalytic subunit (α or R1) (Jordan and Reichard, 1998). The α subunit is encoded by *nrdA* or *nrdE* and the β subunit is encoded by *nrdB* or *nrdF*. These genes form exclusive pairs: *nrdA* is found only with *nrdB* (*nrdAB*), and *nrdE* is found only with *nrdF* (*nrdEF*). Notably, the Class I α subunit is thought to have evolved directly from dimeric Class II RNRs, so they share several catalytic sites (Lundin et al., 2015). The radical initiation mechanism of the β subunit further divides Class I RNRs into five subclasses (a-e) (Blaesi et al., 2018; Cotruvo et al., 2011, 2013; Rose et al., 2018) (Fig. 1b). Subclass Ia uses a diiron cluster activated by O_2_ to oxidize a tyrosine residue, thus forming a stable protein radical (Cotruvo et al., 2011). Subclass Ib also forms a stable radical on a tyrosine residue in the β subunit, but instead uses a dimanganese cluster and is oxidized by superoxide rather than O_2_ (Cotruvo et al., 2013). Subclass Ic is activated by O_2_, but does not form a protein radical (Högbom et al., 2004). Instead, its di-metal cluster (Mn/Fe) is used directly to produce the thiyl radical on the α subunit (Jiang et al., 2007). Like subclass Ic enzymes, subclass Id generates the thiyl radical directly with the use of a di-metal cofactor (Mn_2_) (Rose et al., 2018). However, like subclass Ib, it is unreactive to O_2_ and is activated by superoxide (Cotruvo et al., 2013; Rose et al., 2018). Finally, subclass Ie enzymes are metal-free, instead using a dihydroxyphenylalanine (DOPA) radical as the initiator in an O_2_ dependent reaction (Blaesi et al., 2018). Subclasses Ib and Ie also require a separate flavodoxin activase, NrdI. Class I RNRs are generally presumed to be subclass Ia enzymes unless they can be assigned to another subclass based on sequence homology to a close relative that has been biochemically characterized (Berggren et al., 2017).

While the diversity of RNR biochemistry makes this enzyme an excellent marker for inferring aspects of viral biology, proper annotation of RNR genes is imperative for this purpose. Unfortunately, this same diversity has also fostered high misannotation rates, with one study reporting that 77% of RNRs submitted to GenBank had misannotations (Lundin et al., 2009). Most of those misannotations (88%) were due to RNR sequences being assigned to the wrong class. In response, a specialty database (RNRdb) was created for maintaining a collection of correctly annotated RNRs (Lundin et al., 2009). Even with resources such as the RNRdb, however, the complexity of RNR annotation remains daunting for non-experts. Class I RNRs can be particularly difficult to identify, as their classification relies largely on the annotation of both an α and β subunit.

Our prior work examining the phylogenetic relationships among RNRs from marine virioplankton revealed two large clades of cyanophage RNRs, the first made up of Class I enzymes and the second of Class II RNRs (Sakowski et al., 2014). The hosts of these cyanophage, marine *Synechococcus* and *Prochlorococcus*, carry Class II RNRs. Thus, the presence of such a large cyanophage clade with Class I RNRs was intriguing, and in contradiction to earlier findings that phage tend to carry an RNR gene similar to that of their host cell (Dwivedi et al., 2013). Now, the reanalysis of an RNR from the Class II-carrying cyanophage has revealed that the RNRs in this second clade are, in fact, Class I RNRs that were misannotated as Class II. The reannotation of the RNR from *Prochlorococcus* phage P-SSP7 from Class II to Class I implies that most known cyanophage carry RNRs that are not host-derived, nor dependent on B_12_. Additionally, our analysis suggests that the P-SSP7 RNR may represent a novel Class I RNR subclass.

## 2 Materials and Methods

### 2.1 The Cyano SP Clade

The RNR from *Prochlorococcus* phage P-SSP7 is a member of the ‘Cyano II’ RNR clade, as recognized by Sakowski et al. (Sakowski et al., 2014) in a study of virioplankton RNRs. Based on our analysis, and to avoid confusion with the nomenclature for RNR classes, we have renamed the Cyano II clade to the Cyano SP clade, as RNRs in this clade are exclusively found within the cyanosipho- and cyanopodoviruses (Sakowski et al., 2014). We have also renamed the Cyano I clade to the Cyano M clade, as RNRs in this clade are exclusively seen in cyanomyoviruses. The aforementioned study included ten reference sequences from the (now) Cyano SP clade. Eight of those ten references were used in the current study (Table 1). Cyanophage KBS-S-1A was excluded because its genome has not been fully sequenced and *Synechococcus* phage S-CBP3 was excluded because its RNR was missing a conserved catalytic site. P-SSP7 was chosen as the clade representative because it is the most well-studied phage from this group, has a full genome available, and is the source of the original RNR misannotation.

**Table 1.** Cyano SP clade reference sequences and their hosts.

| **Virus** | **Family** | **Host** | **Host RNR type** |
| --- | --- | --- | --- |
| *Prochlorococcus* phage P-SSP7 | *Podoviridae* | *Prochlorococcus marinus* subsp. pastoris str. CCMP1986 | II – monomeric |
| Cyanophage P-SSP2 | *Podoviridae* | *P. marinus* MIT 9312 | II – monomeric |
| Cyanophage 9515-10a | *Podoviridae* | *P. marinus* MIT 9515 | II – monomeric |
| Cyanophage NATL1A-7 | *Podoviridae* | *P. marinus* str. NATL1A-7 | II – monomeric |
| Cyanophage NATL2A-133 | *Podoviridae* | *P. marinus* str. NATL2A-133 | II – monomeric |
| Cyanophage SS120-1 | *Siphoviridae* | *P. marinus* SS120 | II – monomeric |
| Cyanophage Syn5 | *Podoviridae* | *Synechococcus* str. WH8109 | II – monomeric |
| *Synechococcus* phage S-CBS4 | *Siphoviridae* | *Synechococcus* CB0101 | II – monomeric |

### 2.2 Putative α and β subunit identification

Putative α and β subunit sequences were extracted from the genome of *Prochlorococcus* phage P-SSP7 (genome accession no. NC_006882.2). The putative Class I α subunit is the RNR currently identified in the P-SSP7 genome as ribonucleotide reductase class II (accession no. YP_214197.1) and was downloaded from NCBI in April 2018. As P-SSP7 has no annotated β subunit, candidate β sequences were identified based on length filtering of unannotated protein sequences. While Class I β subunits are typically between 350 and 400 amino acids (Kolberg et al., 2004), we expanded our search range to avoid excluding any potential Class I β subunits. Four candidate, unannotated proteins between 200 and 500 amino acids in length were downloaded for analysis in May 2018. Candidate proteins were searched against the Conserved Domain Database using batch CD-Search (Marchler-Bauer et al., 2017).

The P-SSP7 putative Class I RNR α subunit and four candidate β subunit proteins were imported into Geneious v10.2.4 (https://www.geneious.com) to analyze conserved residues. The putative α subunit peptide sequence was aligned with one representative of each of the known Class I subclasses (Table 2) using the MAFFT v7.388 Geneious plug-in (Katoh and Standley, 2013) on the FFT-NS-ix1000 (iterative refinement method with 1000 iterations) setting with the BLOSUM62 scoring matrix. If necessary, alignments were manually modified to ensure that annotated active sites in the subclass representatives were properly aligned. References have been biochemically characterized and have corresponding crystal structures, where possible. Active sites were annotated for each of the subclass representatives based on literature reports and crystal structures. Residues from the putative P-SSP7 Class I α subunit aligning with active sites in subclass representatives were recorded (Table 3). Candidate Class I β subunit proteins were analyzed individually in the same manner, using the β subunits corresponding to the Class I α subclass representatives (Table 2).

**Table 2.**
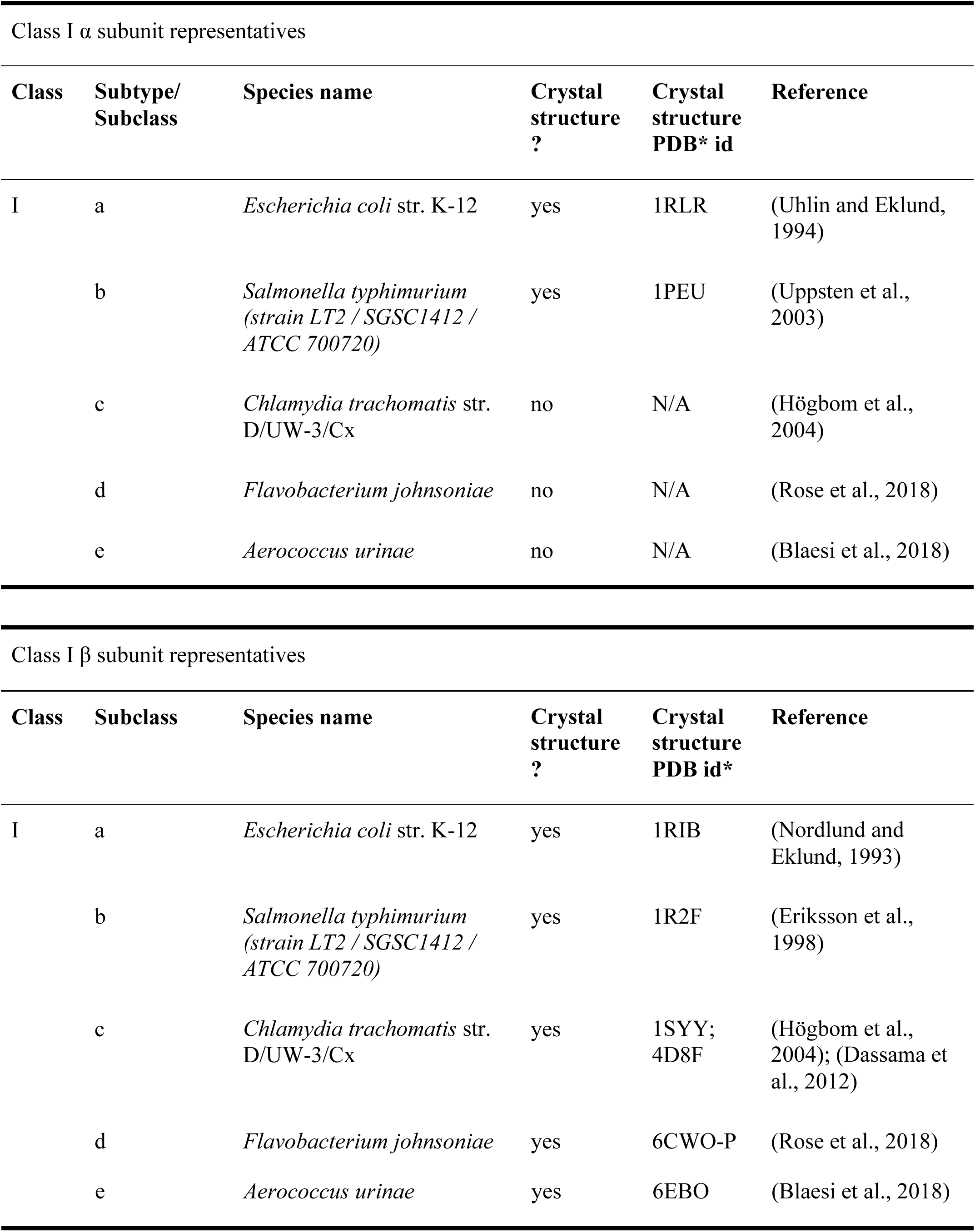
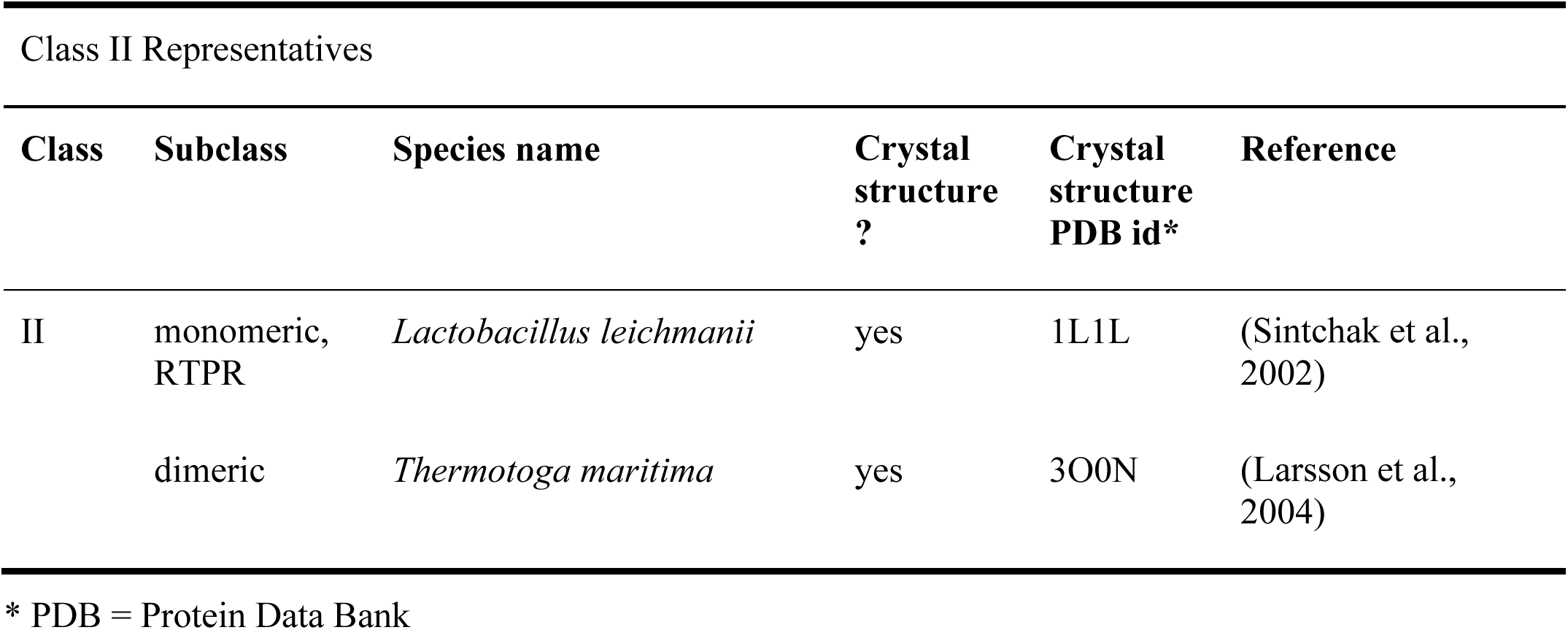
RNR subclass references used for alignment of putative α and candidate β subunits and curation of phylogenetic reference sequences.

**Table 3.** Catalytic residues in Class I RNR α and β subunits and their positions in the putative α and β sequences from *Prochlorococcus* phage P-SSP7. Residues in **bold** were used in reference curation.

| RNR $\alpha$ subunit | | | | |
| --- | --- | --- | --- | --- |
| Residue | Position in P-SSP7 | Position in *E. coli* | Function | Citation |
| **C** | 32 | 225 | active site disulfide bridge | (Lin et al., 1987); (Booker et al., 1994) |
| Q | 105R | 294 | substrate specificity | (Ahmad et al., 2012) |
| R | 110 | 298 | substrate specificity | (Ahmad et al., 2012) |
| N | 187 | 437 | hydrogen bonds | (Kasrayan et al., 2002) |
| **C** | 189 | 439 | thiyl radical - abstracts H | (Mao et al., 1992a); (Mao et al., 1992b) |
| **E** | 191 | 441 | hydrogen bonds | (Persson et al., 1997) |
| **C** | 200 | 462 | active site disulfide bridge | (Mao et al., 1992a) |
| **Y** | 423 | 730 | radical transfer | (Uhlen and Eklund, 1994) |
| **Y** | 424 | 731 | radical transfer | (Uhlen and Eklund, 1994) |
| C | 464 | 754 | radical transfer | (Lin et al., 1987); (Booker et al., 1994) |
| C | 466 | 759 | radical transfer | (Lin et al., 1987); (Booker et al., 1994) |
| RNR $\beta$ subunit | | | | |
| Residue | Position in P-SSP7 | Position in *E. coli* | Function | Citation |
| **W** | 14 | 48 | reduces dioxygen | (Baldwin et al., 2000); (Krebs et al., 2000) |
| Y | 78L | 122 | protein radical | (Larsson and Sjöberg, 1986) |
| F | 122 | 208 | protein radical stability | (Ormö Mats, 1995) |
| F | 127 | 212 | protein radical stability | (Ormö Mats, 1995) |
| R | 145 | 236 | radical transport | (Nordlund and Eklund, 1993);<br>(Eklund et al., 2001) |
| Y | 236 | 356 | radical transport | (Climent et al., 1992);<br>(Rova et al., 1999) |

P-SSP7 candidate β subunit proteins lacking key residues were removed from the analysis. This left a single candidate β subunit protein (accession no. YP_214198.1). Putative active sites identified in the putative β subunit are recorded in Table 3.

### 2.3 Phylogenetic analysis

#### 2.3.1 Phylogenetic reference sequence curation

To create a reference sequence set for phylogenetic analyses, all available Class I α (NrdA and NrdE), Class I β (NrdB and NrdF), and Class II (NrdJ) sequences were downloaded from the RNRdb on August 20, 2018 (Lundin et al., 2009). Sequences were separated into three sets (Class I α, Class I β, and Class II) before sequence curation. Exact and sub-string matches were removed from each set using CD-HIT v4.6 (Fu et al., 2012; Li and Godzik, 2006). Sequences were then divided into smaller groups of similar sequences identified by the RNRdb. RNRdb group assignment is based on phylogenetic clade membership (Berggren et al., 2017; Rozman Grinberg et al., 2018a), so division increased sequence alignment quality. Group names and subclass membership are presented in Table 4. RNRdb sequences were aligned individually by group using the MAFFT v7.388 Geneious plug-in on AUTO setting with the BLOSUM62 scoring matrix. Sequence alignments were visualized and edited in Geneious v10.2.4. Inteins within RNRdb sequences were removed manually after the initial alignment step because they are evolutionarily mobile and confound phylogenetic analyses (Gogarten et al., 2002; Perler et al., 1997). After intein removal, sequences were realigned and those lacking essential catalytic residues were removed, as they are likely non-functional (Sakowski et al., 2014). Other than the two tyrosine residues involved in Class I radical transport (Y730 and Y731, *E. coli*), the same conserved residues were used for Class I α and Class II sequences (Table 3). Both intein removal and catalytic residue identification for all groups were done with guidance from the annotated Class I subclass and Class II representatives (Table 2).

**Table 4.** RNR Class I subclass membership of RNRdb groups.

| Class I subclass | RNRdb groups |
| --- | --- |
| Ia | NrdABe, NrdABg |
| Ia (presumed)* | NrdABh, NrdABk, NrdAm,<br>NrdABn, NrdAq, some<br>NrdABz (NrdABza) |
| Ib | some NrdEF (NrdEFb) |
| Ic | some NrdABz (NrdABzc) |
| Id | NrdABi |
| Ie | some NrdEF (NrdEFe) |

\*The Ia (presumed) subclass includes groups with no biochemically characterized members.

#### 2.3.2 Sequence preparation

Broadly, three categories of phylogenies were constructed from protein sequences: (i) Class I α-only, (ii) Class I β-only, and (iii) Class I α with Class II. All phylogenies included Cyano SP clade members (Table 1). Class I α and Class II proteins share a common ancestor (Lundin et al., 2015), but are phylogenetically unrelated to Class I β proteins. Class I α and Class II proteins also share a common catalytic mechanism and most active sites, but are divergent enough that full-length protein sequences from both classes cannot be presented on the same phylogeny (Lundin et al., 2010). Thus, Class I α and Class II protein sequences in this analysis were trimmed to a previously defined region of interest that excluded regions not shared between the two groups (N437-S625, *E. coli* CQR81730.1) (Sakowski et al., 2014). The Class I α-only phylogeny allowed for greater resolution, as the phylogeny could be based on a longer protein sequence segment, being trimmed only before C225 in *E. coli* (CQR81730.1). Class I β sequences were trimmed to the region between W48 and Y356 (*E. coli*, KXG99827.1). For Class I α-only and Class I β-only phylogenies, sequences were trimmed near the N-terminus to exclude evolutionarily mobile ATP cone domains (Aravind et al., 2000). Class I β sequences were also trimmed near the C-terminus to exclude any fused glutaredoxin domains (Rozman Grinberg et al., 2018b). In all cases, trimming was guided by annotated Class I subclass (a-e) and Class II subtype (mono- or dimeric) representatives (Table 2).

In addition to trimming, sequences were clustered prior to phylogenetic analysis, as each group contained a large number of sequences (Class I α: 15,894 sequences, Class I β: 17,109 sequences, and Class II: 9,147 sequences). To avoid inter-group mixing within individual sequence clusters, sequences were clustered by RNRdb group (Table 4). Clustering of RNRdb sequences was performed at multiple identity thresholds (70%, 75%, and 80%) using CD-HIT v4.7 to ensure that the placement of the Cyano SP clade was not an artifact of the identity threshold, as Cyano SP members have grouped with Class II sequences in the past (Sakowski et al., 2014). Cyano SP sequences were not clustered before phylogenetic analysis. For Class I α-only and β-only phylogenies, sequences were clustered over 80% of the alignment length. For the Class I α with Class II phylogeny, sequences were clustered over 100% of the alignment length due to the short length of the trimmed region.

Two RNRdb groups, NrdABz and NrdEF, contained member sequences belonging to two Class I subclasses (Table 4). In these cases, the Class I β sequences (NrdBz and NrdF) were assigned to subclasses based on active sites. For NrdBz, Class I β subunit enzymes were classified as subclass Ia (NrdBza) by the presence of a Tyr residue in the Tyr radical site (Tyr122 in *E. coli* R2), or as subclass Ic (NrdBzc) by the presence of a Phe, Leu, or Val mutation in the Tyr radical site (Lundin et al., 2009). For NrdF, Class I β subunit enzymes were classified as subclass Ib (NrdFb) or Ie (NrdFe) if carboxylate residues were conserved or missing, respectively, from the second, fourth, and fifth metal-binding sites in relation to the subclass Ib representative (Table 2). Class I α sequences (NrdAz and NrdE), which could not be assigned to subclasses based on primary sequence alone, were assigned to a subclass based on the assignment of their corresponding β subunits. Class I α subunit sequences that were not able to be paired with a β subunit, or that were paired with more than one β subunit, were excluded from further analysis. Excluded Class I α subunit sequences included 1006 NrdAz and 2921 NrdE sequences, or 31% and 45% of total curated NrdAz and NrdE sequences, respectively. The excluded sequences comprised a small percentage of overall RNR diversity (Table S1). Thus, their exclusion is not expected to have affected the phylogenetic analyses (Table S1). All other RNRdb groups exclusively belonged to a single subclass.

#### 2.3.3 Phylogenetic tree construction

For all phylogenetic analyses and clustering identity thresholds, cluster representatives were aligned with correspondingly trimmed α or β subunits from the Cyano SP clade. All alignments were constructed in Geneious using the MAFFT v7.388 plug-in with setting FFT-NS-2 (fast, progressive method) and the BLOSUM62 scoring matrix. Trees were constructed using the FastTree v2.1.5 (Price et al., 2010) Geneious plug-in with default settings. Trees were visualized and customized in Iroki (Moore et al., 2018). Phylogenies inferred from sequences clustered at different identity thresholds can be found in the supplement (Figs. S1-S3).

Finally, a phylogeny was constructed from trimmed Class I α subunit and Class II sequences from only cyanobacteria and cyanophage. No clustering was performed. The phylogeny was constructed as described above from an alignment done using the MAFFT v7.388 plug-in with setting FFT-NS-ix1000 (iterative refinement method with 1000 iterations).

### 2.4 Sequence similarity network

A protein sequence similarity network (SSN) was constructed with the same RNR Class I β subunit sequences used for phylogenetic analysis. The SSN was generated with the Enzyme Similarity Tool (EFI-EST) (Gerlt et al., 2015) as in Rose et al. (E-value: 5, fraction: 1, minimum alignment score: 90) (Rose et al., 2018). As the full network was too large to visualize in Cytoscape (Shannon et al., 2003; Smoot et al., 2011), the 90% identity representative node network was used (i.e., each node in the network contained sequences that shared at least 90% amino acid identity).

## 3 Results

*Prochlorococcus* phage P-SSP7 is a cyanopodovirus that infects the marine cyanobacterium *Prochlorococcus marinus* subsp. *pastoris* str. CCMP1986 (Sullivan et al., 2005). The RNR discovered in P-SSP7 was initially annotated as Class II based on the apparent lack of a Class I β subunit in the phage genome. The RNR from P-SSP7 also lacks an ATP cone region, a domain that is common in Class I α subunits but rare in Class II enzymes (Aravind et al., 2000; Jonna et al., 2015). This was also the first cyanophage RNR of its kind to be annotated, and consequently this gene became the baseline annotation for closely related RNRs. Prior examination of RNRs in viral shotgun metagenomes (viromes) designated the phylogenetic clade containing the RNR from P-SSP7 as the ‘Cyano II’ clade, recognizing that member RNRs (Table 1), exclusively from cyanophage, were annotated as Class II and seemed to fall on the Class II side of the tree (Sakowski et al., 2014). This study also recognized a ‘Cyano I’ clade composed exclusively of cyanomyoviruses that carried Class I RNRs (Sakowski et al., 2014). The Cyano II clade has been renamed to Cyano SP, as the clade is comprised solely of RNRs from cyanosipho- and cyanopodoviruses. The Cyano I clade has been renamed to Cyano M, as it consists of RNRs strictly from cyanomyoviruses.

### 3.1 P-SSP7 Class I α subunit identification

The first indication that the RNR from P-SSP7 was misannotated as a Class II RNR came from the observation of two consecutive tyrosine residues (Y730 and Y731 in *E. coli*) that are present in the C-terminus of Class I α subunits and participate in long-range radical transport between the α and β subunits of Class I RNRs (Greene et al., 2017; Uhlin and Eklund, 1994). These tyrosines are not present in Class II RNRs but are present in the P-SSP7 RNR peptide (Table 2). To confirm the classification of the P-SSP7 RNR as a Class I enzyme, a phylogenetic tree was constructed containing Class I α subunits and Class II sequences from the RNRdb, together with the putative α subunits from the Cyano SP clade (formerly Cyano II) reported in Sakowski et al. (Sakowski et al., 2014) (Fig. 2). Trees were constructed at different clustering identities to ensure that the placement of Cyano SP sequences with a given RNR class was not an artifact of the clustering threshold (Fig. S1). The Cyano SP RNRs grouped with the Class I α subunit sequences in the phylogenies constructed from sequences clustered at 75% and 80% identity, but clustered with Class II sequences in the tree made from sequences clustered 70% identity.

### 3.2 P-SSP7 Class I β subunit identification

While the tyrosine residues within the P-SSP7 RNR are indicative of a Class I RNR, the initial annotation of the P-SSP7 RNR was made primarily because no β subunit gene could be identified within the P-SSP7 genome. Class I RNRs require a β subunit for radical generation. Because the cyanobacterial host of P-SSP7 carries a Class II RNR, the phage would have to carry its own copy of the Class I β subunit gene in order for its α subunit to function. All unannotated proteins in the P-SSP7 genome approximately the length of a Class I β subunit in the P-SSP7 genome were considered RNR β subunit candidates. Four predicted proteins within the genome matched this length criteria. A batch CD-Search (Marchler-Bauer et al., 2017) of the candidate β subunit peptide sequences was unable to identify any conserved domains in any of the sequences. Thus, we aligned the candidate P-SSP7 β subunit sequences with the sequences of biochemically characterized β subunits from each of the known Class I subclasses (Table 2). Only one of the candidate sequences, accession no. YP_214198.1, was found to contain residues experimentally shown to be required for β subunit function (Table 3). The hypothetical protein also resided directly downstream of the α subunit, where the β subunit is typically found (Dwivedi et al., 2013). Thus, YP_214198.1 was identified as the missing P-SSP7 β subunit.

### 3.3 Assignment of P-SSP7 RNR to a Class I subclass

Class I subclasses are based on the mechanism of radical generation utilized by the β subunit. Alignment with representative Class I RNR β subunit sequences found that the P-SSP7 β subunit lacked the tyrosine residue (Y122 in *E. coli* R2) on which the stable protein radical is formed in subclasses Ia, Ib, and Ie (Fig. 1b). The lack of the tyrosine residue seemed to indicate that the P-SSP7 β subunit belonged to subclass Ic, as Ic is the only described subclass that lacks this residue completely (the residue is conserved in Id but does not harbor a radical) (Blaesi et al., 2018; Högbom et al., 2004; Rose et al., 2018). Each subclass has a unique combination of metal-binding residues and uses a different metallocofactor (or does not bind metals at all, in the case of subclass Ie) (Blaesi et al., 2018). The residues in the putative P-SSP7 β subunit aligning with the first sphere of metal-binding residues of the subclass representatives (Table 5) were consistent with Class I RNRs that require metallocofactors (subclasses Ia-Id) and exactly matched subclasses Ic and Id (Blaesi et al., 2018). However, when considering second sphere binding residues, the overall pattern of metal-binding residues in the P-SSP7 β subunit did not match that of any subclass representative (Table 5), nor of any existing RNRdb group (Table 6).

**Table 5.** Metal-binding amino acid residues in each of the β subunit references and P-SSP7.

| Organism | Subclass | First Sphere |  |  |  |  |  | Second Sphere |  |
| --- | --- | --- | --- | --- | --- | --- | --- | --- | --- |
|  |  | 1 | 2 | 3 | 4 | 5 | 6 | 7 | 8 |
| *E. coli* | Ia | D85 | E116 | H119 | E205 | E239 | H242 | S115 | D238 |
| *S. typhimurium* | Ib | D67 | E98 | H101 | E158 | E192 | H195 | M97 | D191 |
| *C. trachomatis* | Ic | E89 | E120 | H123 | E193 | E227 | H230 | E119 | D226 |
| *F. johnsoniae* | Id | E67 | E97 | H100 | E160 | E195 | H198 | C96 | D194 |
| *A. urinae* | Ie | D85 | V116 | H119 | P176 | K210 | H213 | M115 | D209 |
| P-SSP7 | Cyano SP | E42 | E71 | H74 | E117 | E147 | H150 | D70 | D146 |

**Table 6.**
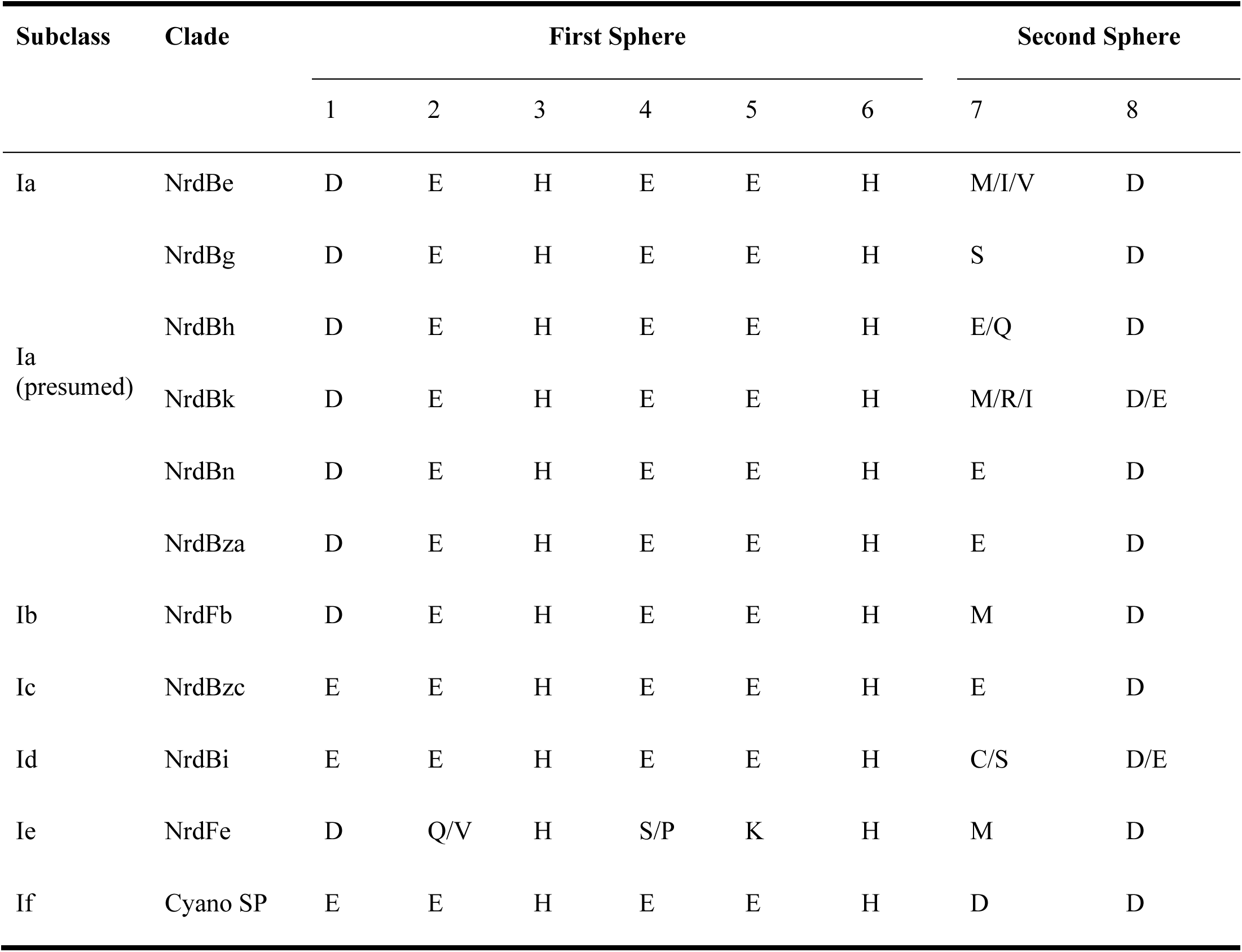
Metal-binding amino acid residues in each of the RNRdb groups and the Cyano SP clade. RNRdb groups are based on phylogenetic clades.

Known Class I subclasses are either monophyletic or contain members that are closely related (Berggren et al., 2017; Rozman Grinberg et al., 2018a). Thus, phylogenetic trees were constructed to confirm proper subclass assignment of the P-SSP7 RNR using Class I β subunit sequences from the RNRdb clustered at 70%, 75%, and 80% and β subunits from the Cyano SP clade members. In a phylogenetic analysis of the 70% identity cluster representative sequences, the P-SSP7 β subunit and Cyano SP homologs were phylogenetically distinct from known RNRs, and did not clearly join with RNRdb groups, instead branching directly off the backbone of the tree (Fig. 3). In the phylogenetic reconstructions at 75% and 80% identity, the Cyano SP grouped remained distinct but branched closely with either the NrdBg group (75% identity, subclass Ia) or the NrdBh group (80% identity, subclass Ia presumed) (Fig. S2). Notably, the Cyano SP β subunits branched away from subclass Ic members (NrdBzc subgroup) in all phylogenies (Fig. S2), making it unlikely that the Cyano SP clade belongs to subclass Ic.

Because Class I subclass assignment was inconclusive based on the β subunit metal-binding residues and phylogenetic analysis, we constructed a protein sequence similarity network (SSN) using the Enzyme Similarity Tool (EFI-EST) (Gerlt et al., 2015) as per Rose et al. (Rose et al., 2018) with the same β subunit sequences used for phylogenetic tree construction (Fig. 4). Most sequences were members of large, distinct subgraphs with sequences exclusively from a single RNRdb group (e.g., NrdBk and NrdBg). However, some RNRdb groups were evenly spread across multiple subgraphs of similar size (e.g., NrdBh and NrdBi), likely indicating a higher level of sequence heterogeneity than other groups. The Cyano SP clade representatives formed exclusive subgraphs not connected to other RNRdb sequences, and were divided into three singleton and one non-singleton cluster, indicating that the clade representatives are divergent even from each other.

Assignment of the Cyano SP RNRs to an existing Class I subclass could not be reliably made based on the analysis of β subunit metal-binding residues, phylogenies, or the protein SSN. Instead, the missing tyrosine radical residue, unique pattern of metal-binding sites, and phylogenetic divergence of the Cyano SP β subunits from RNRdb groups likely indicate that the Cyano SP clade represents a novel Class I subclass.

### 3.4 Origin of the P-SSP7 RNR

Class I α and β subunits tend to evolve in units, producing highly similar phylogenies (Dwivedi et al., 2013; Lundin et al., 2010). Because placement of the Cyano SP β subunits on phylogenetic trees changed with the percent amino acid identity used for clustering RNR sequences (Fig. S2), the Cyano SP α subunits were evaluated for clues to the origin of the RNR in P-SSP7. Class I α-only phylogenies were built from sequences longer than those used for the combined Class I α-Class II phylogenies, allowing greater phylogenetic resolution. Representative RNRdb Class I α subunit sequences from 70%, 75%, and 80% identity clusters were assessed. Regardless of the clustering identity, the Class I α subunit phylogenies showed consistent placement of the Cyano SP clade as an outgroup for the branch that contains RNRdb groups NrdAi (subclass Id) and NrdAk (subclass Ia presumed) (Figs. 5 and S3). Like the Class I β phylogenies, the Cyano SP α subunit clade was distinct and was not surrounded by any RNRdb group. The phylogenetic placement of the Cyano SP Class Iα sequences among RNRdb groups (Fig. 5 & Fig. S3) was different from that seen for the Cyano SP Class I β sequences (Fig. 3 & Fig. S2). Thus, a conclusive placement for the Cyano SP β subunits among RNRdb groups was not possible.

## 4 Discussion

### 4.1 The Cyano SP RNR has adapted to the intracellular environment

The perceived lack of a β subunit gene in the P-SSP7 genome and the lack of an ATP cone domain may have led to the initial misannotation of the P-SSP7 RNR gene as a Class II RNR (Sullivan et al., 2005). Additionally, it seems unusual for a virus to carry a different class of RNR than its host (Dwivedi et al., 2013). Given that cellular organisms carry RNRs that are adapted to their environmental niche (Cotruvo et al., 2011; Reichard, 1993), viruses would also likely benefit from having the same RNR type as their host cell. For example, because marine Cyanobacteria evolved before the Great Oxidation Event (Shestakov and Karbysheva, 2017), they carry Class II RNRs, which do not require oxygen. Widespread iron limitation in the oceans (Moore et al., 2013) and the ability to produce B_12_ (Helliwell et al., 2016) have likely selected against the acquisition of a Class I RNR in marine Cyanobacteria. Thus, given that P-SSP7 would be infecting its host in those same iron limited conditions, and that the acquisition of the host RNR would likely increase its fitness, P-SSP7 might also be expected to carry a Class II RNR.

The preference for a potentially iron-dependent Class I RNR enzyme among cyanophage seems puzzling considering that iron is often the primary limiting nutrient in the oceans, including in regions dominated by *Synechococcus* and *Prochlorococcus* (Browning et al., 2017; Moore et al., 2013). *Synechococcus* and *Prochlorococcus*, hosts infected by phages within the Cyano SP (cyanosipho- and cyanopodoviruses) (Table 1) and Cyano M (cyanomyoviruses) clades, are some of the few B_12_ producers in the oceans (Heal et al., 2016; Helliwell et al., 2016). Therefore, B_12_ availability would seem to be sufficient for viral replication with a B_12_-dependent Class II RNR, while iron availability for phage-infected cells could be too low to support the highly lytic phenotype displayed by many of these phages.

However, carrying a Class I RNR would relieve marine cyanophage of their dependence on the host to produce sufficient levels of B_12_ for deoxyribonucleotide synthesis by a Class II enzyme. Although it is less limiting than iron in ocean waters, B_12_ is likely to be more limiting than iron inside a cyanobacterial cell. In Cyanobacteria, B_12_ is used as a cofactor for two enzymes, the Class II RNR (NrdJ) and methionine synthase MetH (Heal et al., 2016). NrdJ is needed only while the cell is actively replicating, thus, transcription of this gene is closely tied with the cell cycle (Herrick and Sclavi, 2007; Mowa et al., 2009). Similarly, MetH expression is high during early growth of the B_12_-producing cyanobacterium *Synechocystis* but decreases when cells enter the stationary growth phase (Tanioka et al., 2009). Given that NrdJ and MetH are both tied to cellular growth, intracellular B_12_ concentrations are likely highly variable. In addition, cobalt, the metal at the center of B_12_, is required almost exclusively for B_12_ formation and is tightly controlled because of its toxicity to cells (Huertas et al., 2014; Waldron et al., 2009). In contrast, both iron and manganese are required for numerous proteins and molecules within a cyanobacterial cell that are needed throughout the cell cycle (Palenik et al., 2003; Shcolnick and Keren, 2006). Cytoplasmic cyanobacterial iron and manganese quotas have been documented at 10^6^ atoms/cell (Keren et al., 2002, 2004) and a study that aimed to identify and quantify metals in a cyanobacterium found that iron was present in high intracellular concentrations, while cobalt concentrations were below the detection limit (Barnett et al., 2012). Furthermore, some *Prochlorococcus* are able to maintain growth while up-taking just one atom of cobalt per cell per hour (Hawco and Saito, 2018). Therefore, upon infection, a cyanophage would encounter an intracellular pool of iron many fold larger than that of B_12_.

The acquisition of B_12_ from the surrounding environment also seems unlikely. B_12_ is bulky and structurally complex, requiring special transporters which neither *Prochlorococcus*, *Synechococcus*, nor their phages are known to encode (Pérez et al., 2016; Rodionov et al., 2003; Tang et al., 2012). Furthermore, one study showed that while some organisms, such as eukaryotic microalgae, are able to import partial or finished forms of B_12_, *Synechococcus* and likely *Prochlorococcus* are unable to do this (Helliwell et al., 2016). Instead, *Synechococcus* is required to synthesize B_12_ start to finish (Helliwell et al., 2016), likely because both *Prochlorococcus* and *Synechococcus* produce a form of B_12_ that seems to be unique to Cyanobacteria (Heal et al., 2016).

Finally, B_12_ is energetically expensive to synthesize and structurally complex. B_12_ synthesis requires a long pathway made up of roughly twenty different enzymes (Warren et al., 2002). By comparison, some Class I RNR metallocofactors are known to self-assemble (Cotruvo et al., 2011). At most, a metallocofactor may require a flavodoxin (NrdI) for assembly (Blaesi et al., 2018). When considering that carrying a Class I enzyme relieves the phage of relying on a complex host-mediated pathway for a molecule that is not consistently produced throughout the cell cycle, the difference in RNR type between host and phage is not surprising.

The RNR from P-SSP7 also seems to have adapted to the environment inside the host cell in other ways. The P-SSP7 β subunit lacks the tyrosine residue used for radical generation in most Class I RNR subclasses (Fig. 1b). The tyrosine residue harbors a stable protein radical and is a target of nitric oxide (Eiserich et al., 1995; Radi, 2004). Tyrosine-radical scavenging nitric oxide is hypothesized to be present inside *Synechococcus* cells as an intermediate in nitrate reduction (Preimesberger et al., 2017), which is widespread among freshwater and marine *Synechococcus* species and is coupled to photosynthesis (González et al., 2006; Guerrero, 1985; Klotz et al., 2015; Sunda and Huntsman, 2015). Thus, the loss of the tyrosine radical site in the Class I β subunit genes of cyanophage, such as P-SSP7, would enable these phages to avoid RNR inactivation by nitric oxide.

### 4.2 Connections between RNR and cyanophage phenotype

Most Class I RNR α subunits contain an ATP cone region. ATP cones are regulatory sites that essentially act as on/off switches for RNRs (Aravind et al., 2000; Brown and Reichard, 1969). When ATP is bound, the RNR holoenzyme enters a conformational state that allows for function (Eriksson et al., 1997). Once dNTP levels rise high enough, dATP binds the ATP cone and the holoenzyme enters a non-functional conformation (Eriksson et al., 1997; Mathews, 2006). Intriguingly, the Class I α subunits of the Cyano SP clade do not have ATP cones. This is unusual for Class I α subunits and likely represents an evolutionary loss, given that only two Class I α subunit clades (NrdAi/NrdAk and NrdEb/NrdEe) (Fig. 5) lack ATP cones (Aravind et al., 2000; Jonna et al., 2015). In losing the ATP cone domain, the Cyano SP RNRs have lost this regulatory switch. As a consequence, the RNR of cyanopodo- and cyanosiphoviruses cannot be inactivated through dATP binding, thereby leading to unregulated production of deoxyribonucleotides for DNA replication. This phenotype would be beneficial to a fast-replicating lytic phage (Chen et al., 2009).

The highly lytic nature of the Cyano SP clade is also reflected in the biochemistry of the family A DNA polymerase gene (*polA*) carried by some of the members of the clade (Table 1). The amino acid residue at position 762 (*E. coli* numbering) plays a role in shaping the activity and fidelity of Pol I (polA peptide) and is hypothesized to be reflective of phage lifestyle (Schmidt et al., 2014). Prior work found that a mutation from phenylalanine to tyrosine at position 762 produced a 1,000 fold increase in processivity with a concomitant loss of fidelity (Tabor and Richardson, 1987). Three of the member phages within the Cyano SP clade carry a Pol I with a tyrosine at position 762, indicating that Cyano SP members are capable of fast DNA replication. Other members carry *polA* genes that contain a frameshift mutation, preventing identification of the 762 position. Pairing an unregulated RNR, such as the Cyano SP RNR, with a highly processive DNA polymerase would be advantageous for a highly lytic phage. This phenotype is thought to be characteristic of most cyanopodoviruses (Schmidt et al., 2014; Suttle and Chan, 1993; Wang and Chen, 2008). Observations of gene associations such as Tyr762 PolA and Cyano SP clade Class I RNR can thus inform predictions of the possible life history characteristics of unknown viruses.

### 4.3 A novel Class I RNR in cyanophage

Reannotation of the P-SSP7 RNR from Class II to Class I is based primarily on the discovery of a Class I β subunit in the P-SSP7 genome. The P-SSP7 β subunit was identified using conserved residues, as no conserved domains could be identified in the previously hypothetical protein. Our discovery of the Class I β subunit via active sites and genome location demonstrates that some unknown viral proteins (i.e., the viral genetic dark matter) (Krishnamurthy and Wang, 2017) could actually be well known proteins that are simply too divergent for annotation using homology searches or gene model approaches.

The reannotation is also supported by the presence of the consecutive tyrosine residues in the C-terminus of the newly annotated Class I α subunit, which are essential for radical transfer between Class I α and β subunits (Greene et al., 2017; Uhlin and Eklund, 1994) and are not found in Class II RNRs. Additionally, two trees constructed from Class I α and Class II sequences showed the Cyano SP clade (represented by P-SSP7) on the Class I side of the tree (Fig. 2 and Fig. S1b). While the 70% Class I α with Class II tree showed the Cyano SP clade on the Class II side of the tree, we believe this is an artifact of the low identity threshold and short region of interest (Fig. S1a). Protein SSNs constructed from the same sequences used in the Class I α with Class II phylogeny showed the Cyano SP clade as being distinct from both Class I αnd Class II sequences (Fig. S4). Thus, the high divergence of the Cyano SP clade as compared to Class I α and Class II sequences in the RNRdb are likely contributing to the Cyano SP clade grouping with Class II sequences on the 70% tree. Given the presence of the tyrosine residues, the consistent grouping of the Cyano SP clade on the Class I α- only trees, and the presence of the β subunit, we are confident in assigning the Cyano SP clade to Class I. A study of gene transcription in P-SSP7-infected *Prochlorococcus* cultures lends further experimental support for the presence of a Class I RNR in P-SSP7. Both the P-SSP7 Class I RNR α subunit (identified as nrd-020) and the neighboring β subunit (identified as nrd-021) were co-expressed during the second stage of phage infection, during which DNA replication typically takes place (Lindell et al., 2007).

**Figure 2.**
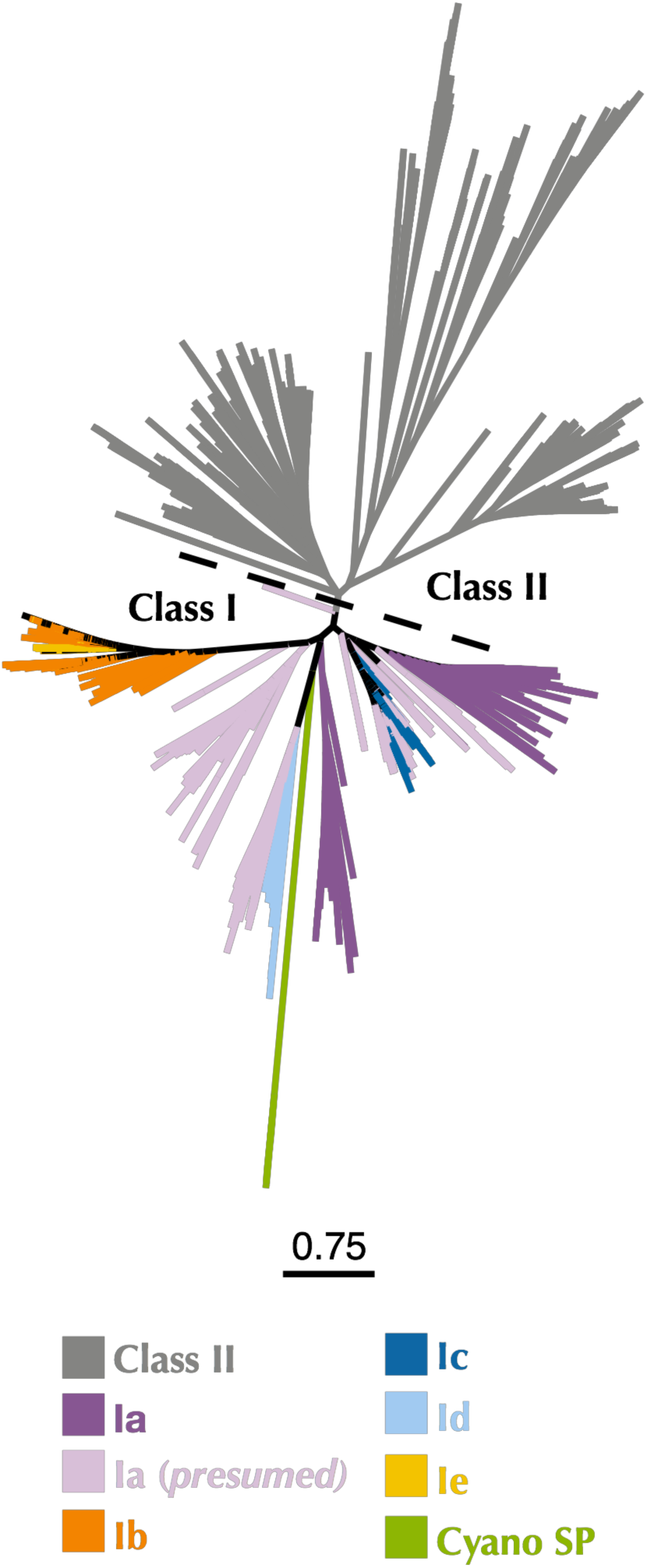
Maximum-likelihood phylogenetic tree of Cyano SP clade α subunits with 80% clustered Class I α and Class II RNRdb sequences trimmed to a region of interest. Gray branches belong to Class II. Colored branches belong to one of the five Class I subclasses, or Cyano SP as indicated in the key. Light purple branches indicate RNRdb groups without characterized members, which are assumed to be subclass Ia enzymes. Trees were constructed using FastTree and visualized and customized in Iroki. Scale bar represents amino acid changes per 100 positions.

Assignment of the P-SSP7 RNR to an existing Class I subclass was inconclusive as the radical-generating β subunit (Cotruvo et al., 2011) could not be clearly assigned based on conserved residues. While the P-SSP7 β subunit contains all of the conserved residues required for function (Table 3), it lacks the tyrosine residue (Y122 in *E.coli*) that harbors the stable protein radical or is conserved in subclasses Ia, Ib, Id, and Ie (Blaesi et al., 2018; Cotruvo et al., 2013; Nordlund and Eklund, 1993) (Fig. 1b). Assignment also could not be made to subclass Ic, the only known subclass lacking the tyrosine residue (Högbom et al., 2004), based on the outcome of phylogenetic (Fig. 3 & Fig. S2) and protein SSN analysis (Fig. 4).

**Figure 3.**
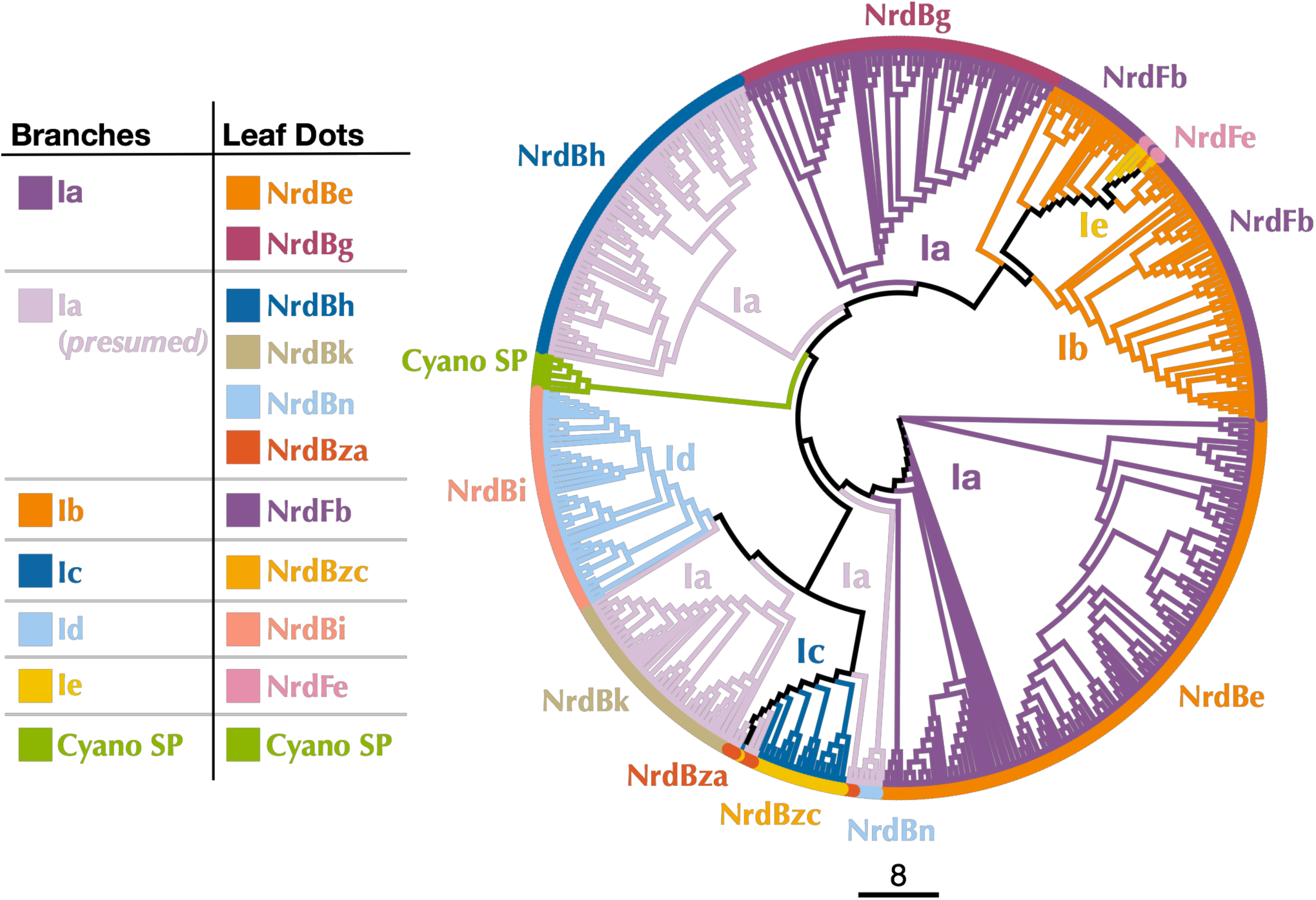
Cladogram of near full-length Cyano SP and 70% clustered RNRdb Class I β subunit sequences. Branch colors indicate Class I subclass and leaf dot colors correspond to RNRdb group. Trees were constructed using FastTree and visualized and customized in Iroki. Scale bar represents amino acid changes per 100 positions.

**Figure 4.**
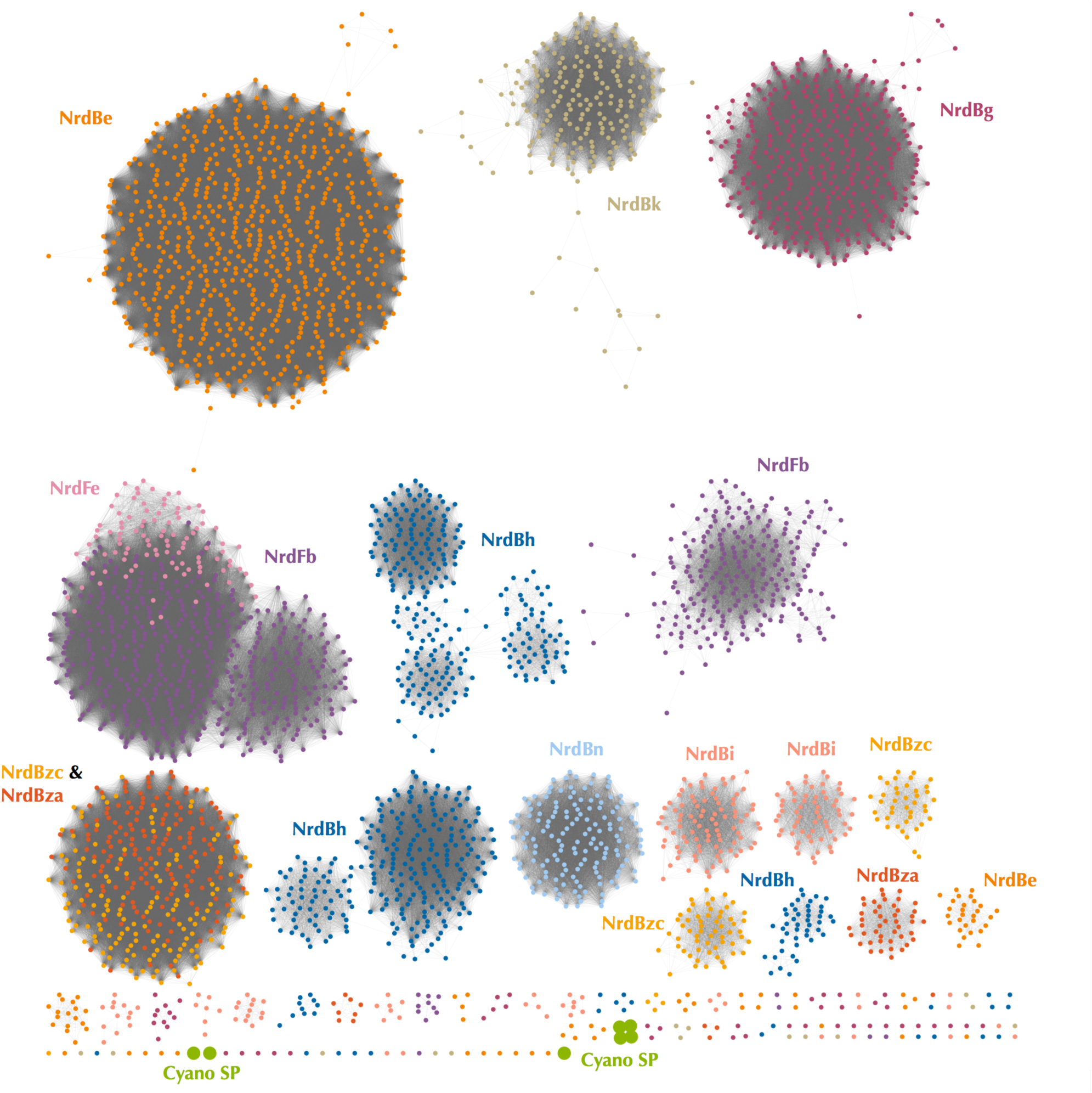
Protein sequence similarity network of the Cyano SP clade and all RNRdb Class I β subunit sequences included in phylogenetic analysis. Nodes represent sequence clusters ≥ 90% similarity. Nodes are colored based on RNRdb group and match leaf dot colors on the cladogram in Fig. 3. Edges connect nodes with minimum alignment score ≥ 90. Network was visualized and customized in Cytoscape.

Additionally, we examined the metal-binding sites in the P-SSP7 β subunit, as metallocofactor identity is used to discriminate between subclasses Ia-Id (Cotruvo et al., 2011; Rose et al., 2018). The metal-binding residues for the P-SSP7 and other Cyano SP clade member β subunits formed a different pattern than is seen in any of the RNRdb groups (Table 6). The combination of the unique metal-binding residues, the lack of a tyrosine residue on which to generate a protein radical, and the phylogenetic distance between the Cyano SP clade and subclass Ic (NrdBzc) sequences, suggest that the P-SSP7 Class I β subunit may constitute a novel subclass of Class I RNRs.

### 4.4 Origin of the P-SSP7 RNR

Because P-SSP7’s host, like most marine *Synechococcus* and *Prochlorococcus*, carries a Class II RNR, we were interested in the origin of the Class I RNR found in P-SSP7. The Class I β subunit phylogenies inconsistently placed the Cyano SP clade. Examination of Class I α subunit trees showed a consistent placement of the Cyano SP clade at the base of the branch harboring the RNRdb groups NrdAk (Ia presumed) and NrdAi (subclass Id) (Figs. 5 and S3). This is perhaps to be expected as, like the NrdAk and NrdAi groups, the Cyano SP Class I α subunits do not contain ATP cone domains, a trait that is rare among Class I α subunits (Jonna et al., 2015).

**Figure 5.**
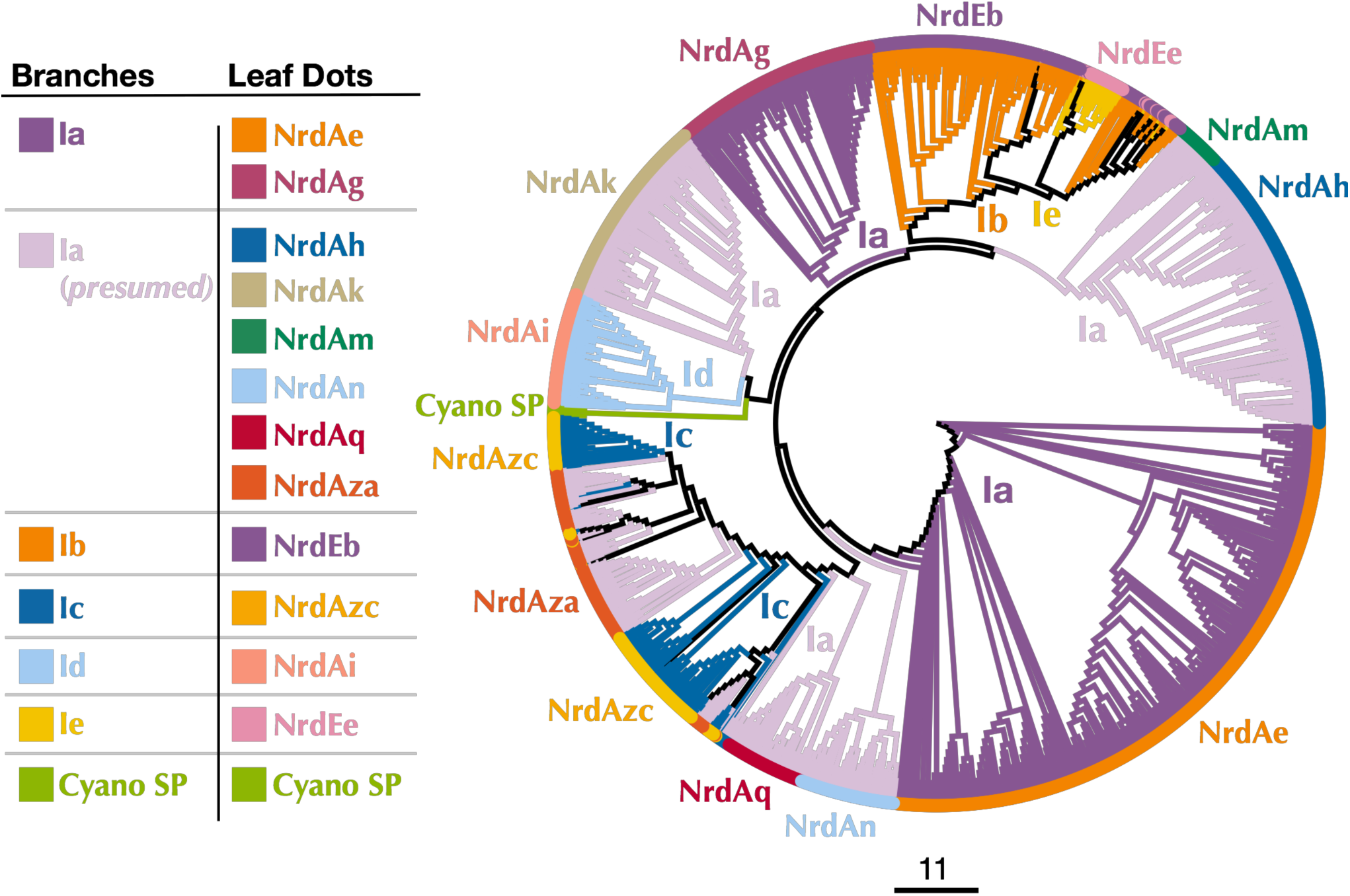
Cladogram of near full-length Cyano SP and RNRdb Class I α subunit sequences clustered at 80%. Branch colors indicate Class I subclass and leaf dot colors correspond to RNRdb group. Colors matching to clades in Fig. 3 indicate α/β subunit pairs. Note there are α subunit clades that do not have corresponding, distinct β subunit clades, as the α subunits have diverged more than the β subunits. NrdAm β subunits belong to β subunit group NrdBh. NrdAq β subunits belong to β subunit subgroup NrdBza. Trees were constructed using FastTree and visualized and customized in Iroki. Scale bar represents amino acid changes per 100 positions.

The observation that the Cyano SP clade does not have the same placement on the Class I β- only and Class I α-only trees is highly unusual. In viruses and cellular organisms, Class I α and β subunits are thought to evolve as units (Dwivedi et al., 2013), producing trees with the same patterns (Lundin et al., 2010). However, viral genomes are known to be highly modular, consisting of genes from multiple sources (Iranzo et al., 2016; Krupovic et al., 2018). It seems possible that an ancestral phage of the Cyano SP clade incorporated the Class I α and β subunits separately. Given that Class I α and β subunits can only perform ribonucleotide reduction as a unit, i.e. both subunits are required for functionality, these acquisitions would have had to occur in quick succession to avoid loss by the phage. Perhaps in support of this hypothesis is that the Cyano SP β subunits sometimes cluster with the NrdBg group (subclass Ia) which harbors the Cyano M clade, while the Cyano SP α subunits consistently cluster with the NrdAi group (subclass Id) that contains the *Synechococcus* phage S-TIM5. These phage groups (i.e. Cyano SP, S-TIM5, and Cyano M) all infect marine *Synechococcus* and *Prochlorococcus*, making the possibility more likely that the Cyano SP RNRs are a mosaic of these cyanomyoviral groups, with the α subunit having been acquired from a cyanophage related to S-TIM5 and the β subunit from a member of the Cyano M clade.

A phylogeny constructed using all Cyanobacteria and cyanophage present in the RNRdb with the Cyano SP clade shows the Cyano SP clade on the Class I side of the tree, distinct from the Class II RNRs (Fig. 6). This phylogeny demonstrates that the majority of known cyanophage carry Class I RNRs. The *Synechococcus* or *Prochlorococcus* hosts of phages in the Cyano M, Cyano SP clades, *Synechococcus* phage S-TIM5, and the Cyanophage P60 clade all carry Class II RNRs (Chen and Lu, 2002; Sabehi et al., 2012; Sakowski et al., 2014). Despite being a myovirus, S-TIM5 does not carry an RNR belonging to the Cyano M clade, likely because it is believed to represent a separate lineage of myoviruses (Sabehi et al., 2012). Interestingly, cyanosipho- and cyanopodoviruses were found in two widely separated clades. Lytic cyanosipho- and cyanopodoviruses within the Cyanophage P60 RNR clade contain a Class II RNR, which is the same type carried by their hosts, whereas cyanosipho- and cyanopodoviruses in the Cyano SP clade contain a Class I RNR. The biological and ecological explanations behind this divergence are a mystery; however, prior work has indicated that cyanopodoviruses can be broadly divided into two clusters, MPP-A and MPP-B, based on whole genome analyses (Huang et al., 2015). Cyanopodoviruses within cluster MPP-B showed greater tendency to carry auxiliary photosynthesis genes, however, no single gene or gene group, including RNR, could clearly distinguish the two clusters. Nevertheless, RNRs belonging to the Cyano SP clade seem to be more common among cyanosipho- and cyanopodoviruses (Huang et al., 2015; Sakowski et al., 2014). Whether carrying a Class II RNR is the ancestral state of cyanosipho- and cyanopodoviruses could not be determined from our phylogenies.

**Figure 6.**
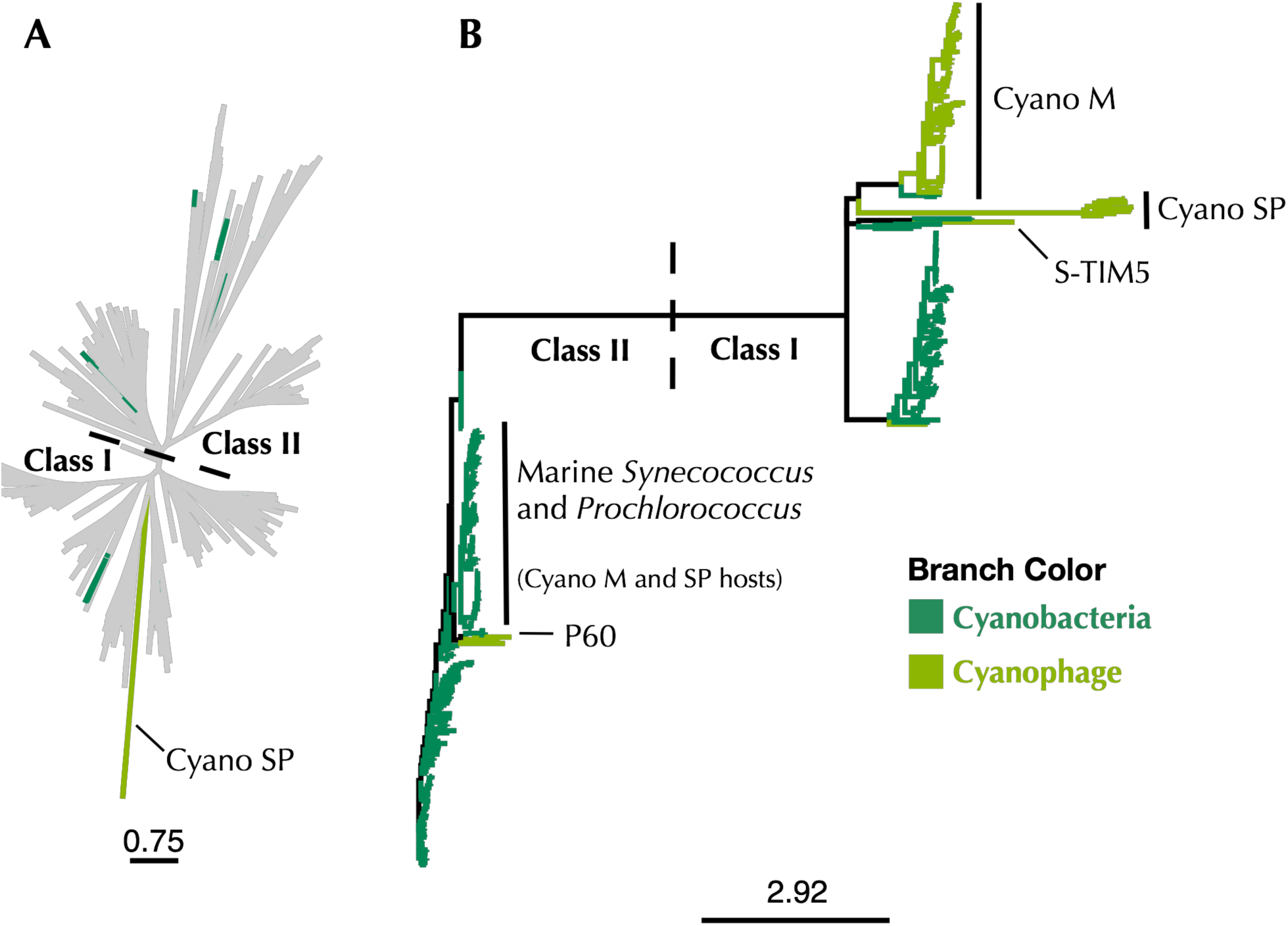
**A)** Maximum-likelihood phylogenetic tree of Cyano SP clade α subunits with 80% clustered Class I α and Class II RNRdb sequences trimmed to a region of interest. **B)** Maximum-likelihood phylogenetic tree of a subset of Class I α subunit sequences limited to Cyanobacteria and cyanophage. In both trees, dark green branches indicate Cyanobacteria and light green branches indicate cyanophage. Trees were constructed using FastTree and visualized and customized in Iroki. Scale bars represent amino acid changes per 100 positions.

The use of marker genes such as RNR in studying viral ecology is important in connecting genomic information to phenotypic traits. However, correct annotation of these genes is essential if accurate information is to be gained. The reannotation also means that most marine cyanophage carry RNRs that did not come from their hosts (Fig. 6), which has implications for our understanding about the acquisition of nucleotide metabolism genes by viruses. That Cyano SP clade members carry Class I RNRs and have lost the tyrosyl radical site in the β subunit is also a reminder that viruses have to adapt to the intracellular environment as well as the extracellular environment. Finally, the discovery of an overlooked β subunit implies that some unknown viral gene space may be composed of known genes that are too divergent for similarity-based annotation methods to detect but can still be identified by other means.

## 5 Conflict of Interest

The authors declare that the research was conducted in the absence of any commercial or financial relationships that could be construed as a potential conflict of interest.

## 6 Author Contributions

AH did the analysis and wrote the manuscript. RM created the sequence similarity networks, assisted with the analysis, and edited the manuscript. KW and SP contributed to study design, data interpretation, and manuscript preparation. All authors read and approved the final manuscript.

## 7 Funding

This work was supported by the National Science Foundation Office of Integrated Activities, grant number 1736030 and the National Science Foundation Division of Biological Infrastructure, grant number 1356374. Computational support by the Univ. of Delaware Center for Bioinformatics and Computational Biology Core Facility was made possible by funding from Delaware INBRE (NIH P20 GM103446) and the Delaware Biotechnology Institute.

## 8 Acknowledgments

We would like to thank Barbra D. Ferrell for critical reading and input on the manuscript.

## 9 Data Availability Statement

The datasets analyzed for this study can be found in the RNRdb (http://rnrdb.pfitmap.org/).

Accession numbers for the Cyano SP clade, including genome accession, can be found in the supplemental material. The supplemental material also contains accession numbers for the annotated RNR subclass representatives.

